# NOISYmputer: genotype imputation in bi-parental populations for noisy low-coverage next-generation sequencing data

**DOI:** 10.1101/658237

**Authors:** Mathias Lorieux, Anestis Gkanogiannis, Christopher Fragoso, Jean-François Rami

## Abstract

**Motivation:** Low-coverage next-generation sequencing (LC-NGS) methods can be used to genotype bi-parental populations. This approach allows the creation of highly saturated genetic maps at reasonable cost, precisely localized recombination breakpoints, and minimize mapping intervals for quantitative-trait locus analysis.

The main issues with these genotyping methods are (1) poor performance at heterozygous loci, (2) a high percentage of missing data, (3) local errors due to erroneous mapping of sequencing reads and reference genome mistakes, and (4) global, technical errors inherent to NGS itself.

Recent methods like Tassel-FSFHap or LB-Impute are excellent at addressing issues 1 and 2, but nonetheless perform poorly when issues 3 and 4 are persistent in a dataset (i.e. “noisy” data). Here, we present an algorithm for imputation of LC-NGS data that eliminates the need of complex pre-filtering of noisy data, accurately types heterozygous chromosomic regions, corrects erroneous data, and imputes missing data. We compare its performance with Tassel-FSFHap, LB-Impute, and Genotype-Corrector using simulated data and three real datasets: a rice single seed descent (SSD) population genotyped by genotyping by sequencing (GBS) by whole genome sequencing (WGS), and a sorghum SSD population genotyped by GBS.

**Availability:** NOISYmputer, a Microsoft Excel-Visual Basic for Applications program that implements the algorithm, is available at mapdisto.free.fr. It runs in Apple macOS and Microsoft Windows operating systems.

Supplementary files: <u>Download link</u>

## Problem

Bi-parental genetic populations can be created from inbred parental lines using various crossing systems, e.g. F_2_ backcross (BC_1_), doubled haploid of F_1_ gametes (DH), F_2_ issued from F_1_ selfing (F_2_), and recombinant inbred lines by single seed descent (SSD). These populations are used to create recombination maps and, if phenotypes are available, to find gene or quantitative-trait locus (QTL) genomic positions.

To do so, each individual of the population under study has to be characterized for its genomic content – or “genotyped” at many loci. This can be done using different molecular biology techniques, including various types of molecular markers. The gold standard for genetic variant discovery is obtained by different next-generation sequencing (NGS) techniques like restriction site-associated DNA sequencing (RADseq)(Davey & Blaxter, 2010), genotyping by sequencing (GBS)(Elshire et al., 2011), and whole-genome sequencing (WGS)(Huang et al., 2009). These techniques provide very large numbers of markers and therefore facilitate the construction of highly saturated genetic maps in mapping populations. This provides accurate locations of recombination breakpoints in each individual, which is important for a number of applications, e.g. studies of local recombination rate, genetic maps comparison, or QTL detection. NGS is still expensive to implement, however, and is commonly applied via reduced representation (RRS-NGS) or low-coverage (LC-NGS) strategies to reduce genotyping costs.

Reducing sequencing costs through minimized per-sample coverage has an important experimental downside: LC-NGS mechanically introduces a series of issues. For example:

– Issue 1: *Low power to detect heterozygosity under low coverage:* For example, if only one sequencing read is generated at a locus, only one of the two alleles is revealed. As each additional read has a 0.5 probability of detecting the second allele, even 3 reads have only 0.25 probability of failing to detect a heterozygous call. Spread over thousands of sites, extensive inaccuracy in heterozygous regions becomes highly problematic.
– Issue 2: *Extensive genotype missingness:* The sparse distribution of reads at low coverage (3X coverage, for example, only implies an *average* of 3 reads per locus) results in a complete lack of reads at some variant loci. Even in plants, which contain more genetic variation than humans, there are only 6-22 variants per 1 Kb, resulting in abundant opportunity for nonreference variant missingness under low coverage.
– Issue 3: *Errors due to erroneous mapping of sequencing reads:* NGS technologies are based on short reads (e.g. 150 bp paired-end Illumina technology). Due to the combinatorial limitation of the sequence contained in short reads, multiple mapping locations may be identified, especially in plant genomes which exhibit much more repetitive content than human genomes. Additionally, in plants, such as rice, it has been shown that structural variation specific to subpopulations may be completely missing in any single reference genome. These assembly errors, omissions, and challenges posed by repetitious regions are sources of erroneous variants. Moreover, outright assembly errors may cause consistent, yet locally encountered genotyping errors.
– Issue 4: *Technical errors inherent to NGS methodology:* Sequencing errors may be globally introduced at a variety of stages in the NGS pipeline, from errors incurred in PCR-dependent library construction to NGS sequencing itself. The initial GBS protocol is known to generate libraries contaminated by chimeric inserts (Heffelfinger et al., 2014). Although rare, these errors may become problematic at low coverage, as additional reads refuting an erroneous call may not be available at a given locus.

Very efficient methods have been recently developed to impute genotypic data derived from LC-NG assays in bi-parental populations. For instance, Tassel-FSFHap (Swarts et al., 2015), LB-Impute (Fragoso, Heffelfinger, Zhao, & Dellaporta, 2016) or Genotype-Corrector (Miao et al., 2018) can all address issues 1 and 2 accurately. Yet, we found in preliminary analyses that these methods can produce inaccurate results when the errors mentioned in issues 3 and 4
 – thereafter called “noisy data” – are too frequent. Thus, these methods might require complex bioinformatic pipelines to filter out low-quality markers before and after imputation. Even then, low-quality markers might not be detected easily and could alter dramatically the quality of the imputation and the final genetic map.

Here, we present an algorithm for imputation of LC-NGS data that eliminates the need of complex pre-filtering of noisy data, accurately finds heterozygous chromosomic regions, corrects erroneous data, and imputes missing data. We test its accuracy using simulated data, and we compare its performance with Tassel-FSFHap, LB-Impute and Genotype-Corrector using three real datasets: (1) a rice SSD population genotyped by GBS and (2) by WGS and (3) a sorghum SSD population genotyped by GBS. The algorithm is implemented in NOISYmputer, a user-friendly computer program (see “Availability and system requirements” section).

## Algorithm

### Pre-requisites

Population genotypes encoded in a Variant Call Format (VCF) file with chromosome coordinates – i.e., we consider organisms that have a reference genome assembly (RefSeq) only. Ideally, the VCF should contain only bi-allelic singlenucleotide polymorphisms (SNP). We haven’t tested VCF that contain small indels. Parental lines need to be included in the VCF file. The VCF file can be compressed (gzip “.gz” format).

### Data preparation

– Convert the VCF to an ‘ABH-’ matrix of genotypes (plain text file) encoded as “A” (homozygous parent 1), “B” (homozygous parent 2, “H” (heterozygous) and “-” (missing data), using parental information. If parental genotypes contain many missing data, it is desirable to pre-impute them from the population. LB-Impute – either stand-alone or from within MapDisto (Heffelfinger, Fragoso, & Lorieux, 2017). – and Tassel-FSFHap can both perform this task.
– Split the ABH-matrix into individual chromosomes.

NOISYmputer has commands to execute these tasks automatically. The VCF conversion to ABH-format is executed by the Java MapDisto Addons. The matrix-splitting command has an option to compress individual chromosomes files. It uses the 7-zip utility, available for free at 7-zip.org.

### Imputation algorithm

We briefly describe the imputation algorithm, which is applied separately to each chromosome. Parameter values for different population types and number of SNPs are suggested in the user manual embedded in the NOISYmputer program.

1. **Filter out SNPs** that contain high percentage of heterozygosity (SSD and DH populations only), high percentage of missing data, and very low minimum allele frequency (MAF).
2. **Filter out redundant loci**, that is, closely linked loci that contain exactly the same genotypic information.
3. **Correct high-likelihood erroneous heterozygote calls** with a very high threshold, using genotypic frequency information of *n* SNPs around each tested locus (default *n* is 14). We use a very high threshold to avoid eliminating real heterozygous calls. *Used for SSD and DH populations only*.
4. **Fill missing data** between identical genotypes only with the same genotype. *This step is necessary to calculate meaningful chi-squared in the next step, but an option is provided to impute later from raw-filtered data that helps eliminating rare errors introduced by this step (for instance, if two linked parental SNPs were wrongly called)*.
5. **Filter out incoherent loci**. *This is a crucial step for noisy data filtering. It allows to find SNPs that do not segregate the same way as their immediate environment, indicating a probable mapping error. As segregation distortion is a frequent phenomenon, we cannot use an overall rule – like comparing to a 1:1 segregation*. Define a window of *n* SNPs around each tested locus. For each window/SNP couple, calculate the A, B and H frequencies across the population, and compare the SNP segregation to the window segregation using a chi-squared test, where expected counts are the observed frequencies in the window multiplied by the population size. Filter out SNPs with a chi-squared statistic that exceeds a define threshold. Parental genotypes can be miscalled, resulting in a wrong encoding of the SNP (BHA-instead of ABH-). To avoid eliminating such SNPs, we also calculate the opposite chi-squared test by swapping f(A) and f(B). If not significant, we retain the SNP and recode it to proper ABH-.
6. **Correct obvious singletons**. *A singleton is an isolated data surrounded by a different allele, for instance the “*B*” in “*AABAA*” is a singleton*. Replace a data at locus *x* if the genotypes of the two previous (*x* – 1 and *x* –2) and the two next loci (*x* + 1 and *x* + 2) are identical and different to the genotype at locus *x*. For instance, “AABAA” is replaced by “AAAAA”. This step is repeated before Step 9 (Impute data-step 2) and Step 11 (Impute data-step 3).
7. **Impute data-step 1**. *Because LC-NGS generates very poor information in heterozygous regions, we use the number of transitions instead of the H frequency to call heterozygotes. A transition is any allelic change between two adjacent SNPs, except missing data*. Define a window of *n* SNPs around each tested locus. In each individual or line, count the number of transitions *T* in the window and calculate the transition rate *TR = T/ n*. Calculate frequencies of A, B and H in the window. Call a homozygous if its corresponding allelic frequency is superior to a defined minimum. Call a heterozygous if a homozygous has not been called and the *TR* is superior to a defined minimum.
8. **Correct obvious singletons** (identical to step 6).
9. **Impute data-step 2**. *Now that we have identified heterozygous regions, we can use frequencies to refine the allele calling*. Define a smaller window of *m* SNPs *(m < n)* around each tested locus. In each individual or line, calculate frequencies of A, B and H in the window. Call a homozygous if its corresponding allelic frequency is superior to a defined minimum. Call a heterozygous if its corresponding allelic frequency is superior to a defined minimum.
10. **Correct obvious singletons** (identical to step 6).
11. **Impute data-step 3**. *We now have a fair representation of A, B and H regions. We further refine the calling using a simple rule based on the maximum allelic frequency*. Define a smaller window of *o* SNPs (*o* < *m*) around each tested locus. In each individual or line, calculate frequencies of A, B and H in the window. Call the maximum frequency.
12. **Extend imputation to chromosome ends**. *Because a full window cannot be defined around loci that are located at a distal position* — *near chromosome ends* —, *these (few) positions remain unimputed*. Call genotypes for terminal SNPs based on the immediately adjacent SNPs that have non-missing data.
13. **Correct singletons in recombination breakpoints**. *The previous steps can generate some rare singletons near the recombination breakpoints*. Correct a singleton at locus *x* using the closest genotype based on physical positions of loci *x, x* – 1 and *x* – 2.
14. **Correct improbable small chromosome chunks**. *The previous steps can generate some improbable chromosome short haplotypes – or chunks*. Identify each short chunk composed of identical alleles, embedded in a homogeneous genomic environment that has a different allele. The genomic environment size is a multiplier of the chunk size (in number of positions) that can be parametrized. Calculate the recombination rate *r_E_* across the population between the left and right bounds of the environment. Calculate the recombination rate *r_C_* across the population between the left and right bounds of the chunk. Replace the chunk allele by its environment allele if the *r_E_ < min_r_E_* or *r_C_ > max_r_C_*.
15. **Collapse the imputed matrix (optional)**. *We do not need the complete imputed matrix, as the only informative SNPs are the ones that mark recombination breakpoints*. Retain the SNPs that mark recombination breakpoints (at one or both sides of the breakpoint).

## Accuracy tests on simulated data

### Method for accuracy estimation

To test NOISYmputer’s accuracy, we used simulated datasets generated as follows: a chromosome of 180 cM containing 10001 SNPs was simulated for 200 SSD lines in MapDisto 2. The calculated map in MapDisto was 172.7 cM. The resulting matrix of genotypes was then imported into NOISYmputer and different types of noise were added to the data using NOISYmputer simulation code: (1) variable missing data rate (MDR), (2) variable MDR and segregation distortion on clustered SNPs, and (3) combinations of missing data rate, A ➜ B or B ➜ A calling error rate, and A or B ➜ H calling error rate. Segregation distortion on clustered SNPs is observed when reads from different genomic regions — *e.g*., gene duplication or other structural variation events — map to the same location on the reference sequence.

We then imputed the noisy data and calculated accuracy as a simple similarity score — or concordance — between the imputed matrix and the initial matrix without noise. Three replicates were performed for each case. Details on simulation and imputation parameters are provided in Table S2 of the Supplementary Tables.

### Results

When varying the missing data rate only, accuracy ranged from 99.63 % to 99.95 % (Figure 1 A, Situation 1, left). When simulating clusters of SNPs showing segregation distortion, similar results were observed with the exception of one datapoint at 98.21 % (Figure 1 A, Situation 2, left). The latter was caused by two closely linked clusters of distorted SNPs and could have been solved using a wider window in step 5 of the algorithm.

**Figure 1.**
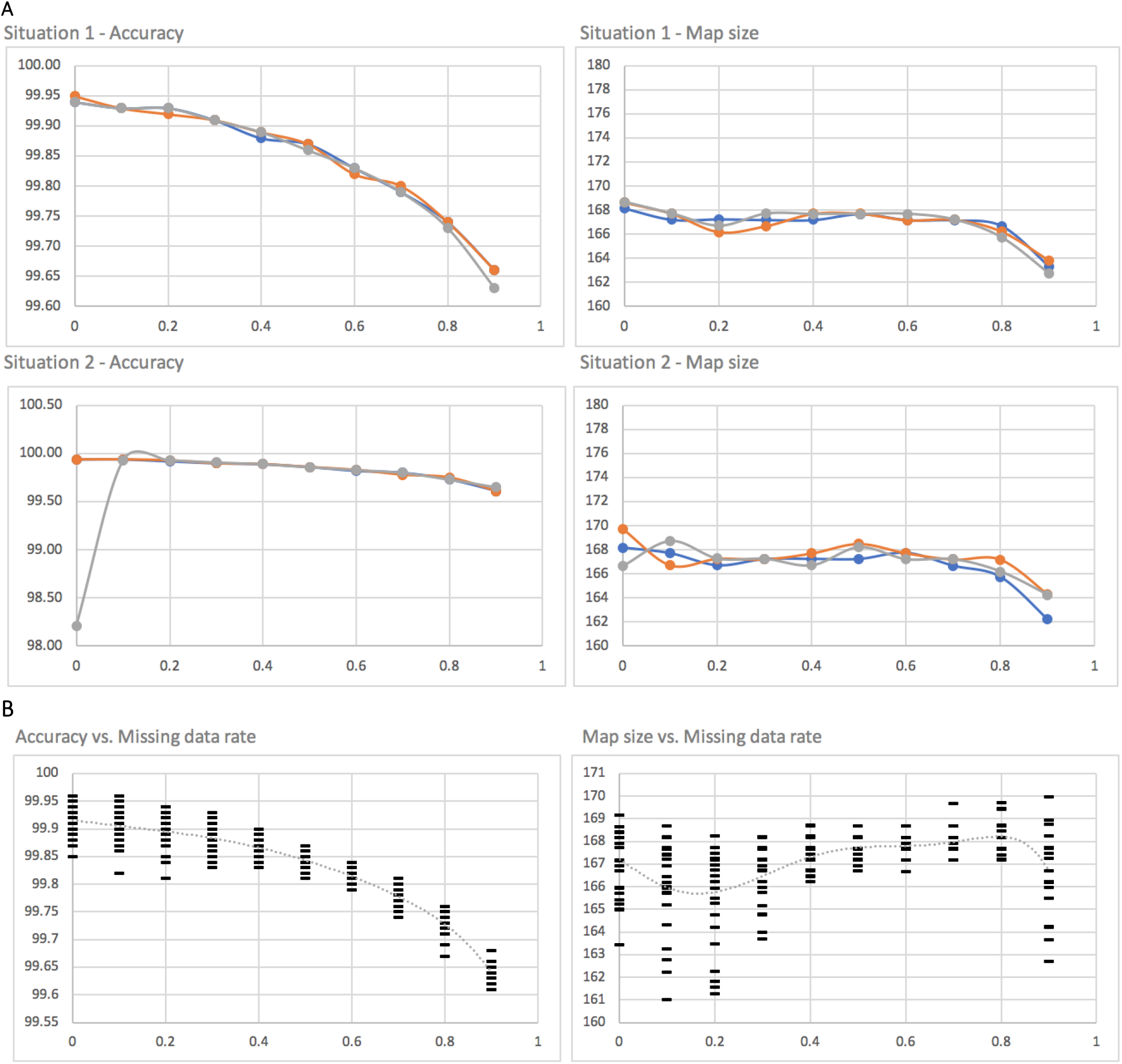
Imputation accuracy (left) and final map size (right) for simulated data, using NOISYmputer. The horizontal axis is the missing data rate in all graphs. A, Situation 1: varying missing data rate (three replicates). A, Situation 2: varying missing data rate and clusters of SNPs showing segregation distortion (three replicates). B: varying missing data rate (MDR). For each MDR value we apply a mix of varying A ➜ B or B ➜ A calling error rate, and A or B ➜ H calling error rate, both ranging from 0.005 to 0.025. Map size is in centiMorgans.

When applying combinations of MDR, A ➜ B or B ➜ A calling error rate, and A or B ➜ H calling error rate, accuracy ranged from 99.61 % to 99.96 % (Figure 1 B, left).

In all cases, map size calculated from imputed data was slightly inferior to the real map size (172.7 cM) (Figure 1 A and B, right). This was likely to be due to too stringent parameters in step 14 that eliminated small improbable chunks.

Overall, accuracy was very high in all situations and comparable to values obtained by authors of Tassel-FSFHap, LB-Impute, and higher than Genotype-Corrector accuracy when MDR exceeded 0.4.

### Application to real data, validation and comparison with other methods

To test the efficiency of NOISYmputer with real data, we compared it with FSFHap, LB-Impute and Genotype-Corrector using and three datasets: one high-quality dataset, and two noisy datasets. We also considered ABHgenotypeR (Furuta, Ashikari, Jena, Doi, & Reuscher, 2017) in early tests but did not go on with full testing as the algorithm showed very poor performance with noisy data. PlantImpute (Hickey, Gorjanc, Varshney, & Nettelblad, 2015) and AlphaPlantImpute (Gonen et al., 2018) were also designed for imputation of bi-parental genetic data, but they only handle DNA chip-based SNP data in their present form. The authors plan to develop a version adapted to GBS data and it will be interesting to test it when available (Gonen et al., 2018).

### Data description

The high-quality real “Rice_GBS” dataset was composed of GBS data (~0.67 X coverage, “Yale protocol” (Heffelfinger et al., 2014)) for a population of 187 rice SSD lines from the cross [IR64 x Azucena], obtained at the IRD, Montpellier, France (Fragoso et al., 2017).

The two noisy real datasets were (1) “Rice_WGS”, composed of WGS data (~1.55 X coverage, “Genoscope protocol” (Karine Labadie, pers. comm)) for 212 SSD lines from the same [IR64 x Azucena] cross, and (2) “Sorghum_GBS”, composed of GBS data (~0.77 X coverage, “Cornell protocol” (Elshire et al., 2011)) for a population of 150 sorghum SSD lines from the cross [SSM1611 x SSM249] kindly provided to us by Dr. Mbaye Ndoye Sall (ISRA/CERAAS, Thies, Senegal). Note that a 1.55X WGS represents lower coverage than a 0.67X GBS, as genome is not reduced in WGS, resulting in fewer reads per site.

Details of how the real data were generated are summarized in Supplementary Table S1 of the Supplementary Data.

### Methods for algorithm comparison

In real data, we do not know the true state at each locus, so we cannot estimate imputation accuracy directly. One alternative way is to calculate the final genetic map and to compare it with existing, high-quality maps. A correctly imputed dataset should generate a map size coherent with maps calculated from high-quality marker data. An imputed dataset that contains substantial rate of genotyping errors will generate *map expansion*, that is, a genetic map longer – in centimorgans (cM) – than its expected size, due to falsely imputed recombination breakpoints.

Map expansion was calculated using the smallest map as a baseline. Maps were calculated for chromosome 1 of each dataset, after full imputation of the few remaining missing data using the NOISYmputer “Full imputation” command. Marker orders were left as determined by the RefSeq. Recombination fractions were obtained with the “Calculate final map” command in NOISYmputer, which is equivalent to the MapDisto 2 two-point method that can use heterozygotes to calculate recombination frequencies in SSD populations (Heffelfinger et al., 2017). We could have also used multipoint mapping as implemented in Mapmaker/EXP (Lander et al., 1987) or R/qtl (Broman, Wu, Sen, & Churchill, 2003), but these algorithms cannot use residual heterozygosity to calculate recombination frequencies in RIL-SSD populations. Note that two-point estimation of recombination fractions on fully imputed data is equivalent to multipoint estimation as implemented in Mapmaker/EXP or R/qtl.

To test NOISYmputer and other methods’ accuracy on real data, we used the high-quality Rice_GBS dataset. We compared map sizes to the map obtained with LB-Impute combined with BP-Impute and a custom R script that selects the most probable genotype, as extensive testing on this dataset showed that this method provided accurate imputation (Fragoso et al., 2016, 2017).

To test performance of the four algorithms with noisy data, we (1) compared the maps they generated for the Rice_WGS dataset to the map obtained for the Rice_GBS dataset and (2) compared map sizes for Sorghum_GBS datasets to the consensus sorghum map (Mace et al., 2009).

Additionally, in order to avoid any bias in map expansion estimation of chromosome 1 only, we calculated the total map size for all chromosomes with NOISYmputer.

Finally, we also tested NOISYmputer’s ability to estimate residual heterozygosity in RILs-SSD, which was simply calculated as the percentage of heterozygous calls over all chromosomes, and compared with the expected residual heterozygosity, ***H_r_*** = 0.5^(*g*–1)^, where *g* is the number of selfing generations.

For Genotype-Corrector tests, we observed that its ‘vcf2map’ command requires a VCF with phased data (A=ref, B= alt) to generate proper ABH-matrices. That is, ‘vcf2map’ doesn’t seem to have an option to use parental information for phasing. Thus, we used the MapDisto Addons VCF converter instead, available in NOISYmputer.

NOISYmputer was run in 64-bit Windows 10 (Apple Bootcamp), and Tassel-FSFHap and LB-Impute were run in macOS with Java 1.8 JRE, on a portable computer equipped with a “Kaby Lake” Intel Core i5 CPU 7267U@3.1 GHz and 16 GB of RAM. Genotype-Corrector was run in Red Hat Enterprise Linux Server release 5.8 (Tikanga), on a Dell PowerEdge M610 computer (server) equipped with an Intel Xeon CPU E5620@2.40GHz and 198 GB of RAM.

Details on parameters used for the three imputation methods are provided in Supplementary Data, section “Imputation commands and parameters”.

## Results

### High-quality real data (Rice_GBS dataset)

All four algorithms performed well when applied to the Rice_GBS high-quality data (Table 1 and Table S3, Rice_GBS entry). NOISYmputer produced the shortest map (183.4 cM), with no apparent imputation incoherence (Figure S1). LB-Impute introduced incorrect heterozygote calls at recombination breakpoints (Figure S2), leading to a map size of 192.1 cM and map expansion of 4.7 % but this can be corrected with BP-Impute (Fragoso et al., 2017). FSFHap produced very similar results compared with NOISYmputer, with a map size of 184.3 cM and a map expansion of 0.5% (Figure S3). Genotype-Corrector introduced incorrectly heterozygote calls at random sites, leading to a 7% map increase (Figure S4), but this could be easily corrected with further and simple chunk correction. We obtained similar results for the other 11 chromosomes.

**Table 1.** Summary of imputation results for chromosome 1 of one high-quality and two noisy datasets, using NOISYmputer, LB-Impute, Tassel-FSFHap and Genotype-Corrector (G-Corrector). (*): SNP number after pre-filtering; (**): SNP number after ABH conversion. (***): LB-Impute could not handle the Rice_WGS dataset. See Table S3 for more details

| *Dataset* | *Algorithm* | *SNPs-*<br>*initial* | *SNPs-*<br>*imputed* | *SNPs-*<br>*collapsed* | *CPU time*<br>*(minutes)* | *SNPs /*<br>*sec.* | *Map size*<br>*(cM)* | *Map*<br>*expansion*<br>*(ratio)* |
| --- | --- | --- | --- | --- | --- | --- | --- | --- |
| **Rice_GBS** | NOISYmputer | 10,850 | 7,489 | 977 | 1.3 | 99.2 | 183.4 | 1 |
|  | LB-Impute | 10,850 | 10,850 | 976 | 121.7 | 1.5 | 192.1 | 1.047 |
|  | FSFHap | 10,850 | 10,850 | 985 | 0.6 | 285.5 | 184.3 | 1.005 |
|  | G-Corrector | 10,850 | 3,161 | 1,811 | 36.2 | 1.5 | 196.2 | 1.07 |
| **Rice_WGS** | NOISYmputer | 210,160 | 91,332 | 1,379 | 8.3 | 419.7 | 181.3 | 1 |
|  | LB-Impute*** | - | - | - | - | - | - | - |
|  | FSFHap | 203,614* | 203,512** | 55,763 | 9.3 | 364.2 | 14,235.4 | 78.5 |
|  | G-Corrector | 210,160 | 40,839 | 5,817 | 760 | 0.9 | 1,231.7 | 6.8 |
| **Sorghum_GBS** | NOISYmputer | 11,516 | 3,837 | 846 | 1.0 | 191.6 | 181.7 | 1 |
|  | LB-Impute | 14,496 | 7,757 | 2,654 | 104.0 | 1.3 | 9,687.0 | 53.4 |
|  | FSFHap | 14,496 | 7,731** | 967 | 0.5 | 258.2 | 1,058.4 | 4.8 |
|  | G-Corrector | 11,516 | 3,595 | 707 | 30.9 | 1.9 | 672.3 | 3.7 |

The total map size calculated with NOISYmputer, at 1,433.8 cM, was very close to the one calculated with LB-Impute combined with BP-Impute and the custom R script, which was 1,439.9 cM (Fragoso et al., 2017). The residual heterozygosity over all chromosomes calculated with NOISYmputer, at 0.037 %, matched very well to the expected heterozygosity (0.049 %).

### Noisy real data (Sorghum_GBS dataset)

With noisy data, we observed important differences in map sizes between the different methods (Table 1 and Table S3, Sorghum_GBS entry). NOISYmputer generated the shortest map, at 181.3 cM, coherent with the consensus sorghum map size of 191.8 cM (Mace et al., 2009) (Figure S5). LB-Impute inaccurately imputed many singletons, resulting in very severe map expansion (53.4X) (Figure S6). FSFHap also inaccurately imputed many singletons, although in lesser extent than LB-Impute, resulting in severe map expansion (5.8X) (Figure S7). Genotype-Corrector was unable to eliminate some noisy data and converted them to heterozygous calls, resulting in severe map expansion (3.7X) (Figure S8).

The total map size calculated with NOISYmputer, at 1,330.4 cM, matched well with the framework sorghum map size (1,355.4 cM). The residual heterozygosity calculated with NOISYmputer, at 1.43%, matched very well to the expected heterozygosity (1.56 %).

### Very noisy real data (Rice_WGS dataset)

The noisiest dataset, Rice_WGS, also revealed the greatest differences in final map sizes between the different algorithm methods (Table 1 and Table S3, Rice_WGS entry). NOISYmputer generated the shortest map, at 181.3 cM, coherent with the Rice_GBS map size of 183.4 cM (Figure S9). The inability of FSFHap to eliminate noisy data, especially clustered distorted SNPs, and the inaccurate imputation of many small heterozygous chromosome chunks, conducted to severe map expansion (78.5 X) (Figure S10). Genotype-Corrector was also unable to eliminate some noisy data and converted them to heterozygous calls, resulting in severe map expansion (6.8 X). It also left many loci unimputed (Figure S11).

The total map size calculated with NOISYmputer, at 1,460.3 cM, was very close to the expected one (1,439.9 cM). The residual heterozygosity calculated with NOISYmputer, at 0.046 %, matched very well to the expected heterozygosity (0.049 %).

### Speed

The four tested algorithms used different programming languages, so comparing them for speed is somewhat unfair. Also, LB-Impute imputes VCF files, while NOISYmputer, FSFHap and Genotype-Corrector impute Hapmap-like files, which induces another unfairness towards LB-Impute. For GBS data, we found that NOISYmputer was greatly faster than LB-Impute (~96 X) and Genotype-Corrector (~28X), and approximately half as fast as FSFHap. For WGS data, NOISYmputer was slightly faster than FSFHap, and ~90X faster than Genotype-Corrector (Table 1).

## Conclusion

Given the results obtained with simulated and real data, we recommend the use of NOISYmputer to impute noisy or high-quality GBS or WGS data. LB-Impute or Tassel-FSFHap can also be used for imputation of high-quality data, with a preference for LB-Impute in case of very low-coverage (< 0.4 X), as discussed in (Fragoso et al., 2016).

We found Genotype-Corrector to be the least accurate of the four methods, even for high-quality GBS data. Genotype-Corrector, according to its developers, does not perform well when missing data rate exceeds 40%, which is the case we treat here. This likely explains its poor performance in our study. Also, Genotype-Corrector filters out markers with segregation distortion, possibly resulting in several gaps in the genetic map of crosses with frequently distorted regions. Furthermore, its authors do not seem to have tested it against FSFHap or LB-Impute, which are specifically designed for bi-parental population imputation (Miao et al., 2018). Instead, they compared it with Beagle, designed for diversity panels and LinkImpute, designed for non-model organisms with no RefSeq available. This might explain why they found Genotype-Corrector to perform better than alternative methods. Finally, we estimated its total run time at 4.35 days for the complete Rice_WGS dataset, which would make it impractical for very large datasets.

One way to even increase imputation accuracy could be to use combinations of NOISYmputer and LB-Impute or FSFHap, which leverage very efficient hidden Markov chain-based methods. For instance, one could use an intermediate output *(e.g*., after the incoherence filter) of NOISYmputer as input to LB-Impute or FSFHap.

We are currently in the process of converting NOISYmputer’s code to Java, in order to further improve its speed. This will allow us to explore wider parameter spaces and, for instance, find sets of parameter values that lead to convergence of an objective function, *e.g*. expected map size or overall heterozygosity.

Although previous methods have made significant advances in addressing the challenges listed above, the noisiness of imputed datasets are still evident by expanded genetic maps, excess heterozygosity, and probabilistically unlikely recombination events contained within a short physical interval. Here, we introduce an algorithm that, in a series of steps, addresses each source of error to create higher-quality datasets for improved trait mapping and genomics-assisted breeding. Although we recognize that, in the near future, new machine learning approaches will likely replace traditional imputation algorithms to extract features representing sources of error, our algorithm represents a step to systematically address all sources of NGS genotyping error, and the corrections made here will be recapitulated in future algorithm development.

## Availability and system requirements

The algorithm is implemented in Visual Basic for Applications (VBA). NOISYmputer, a Microsoft Excel application that implements the algorithm, is available at no charge for academic use (see information at mapdisto.free.fr/NOISYmputer). Researchers who intend to use NOISYmputer for commercial purpose should contact us. NOISYmputer runs in Apple OS X or macOS and Microsoft Windows operating systems. The VCF converter needs Java 1.8. In Windows, 64-bit versions of both the OS and Excel should be preferred for very large datasets. The version for Apple macOS runs better in the 32-bit Excel 2011, as the 2016 and later versions of Excel (2019 or 365) implement a slow, sometimes buggy VBA interpreter.

## Acknowledgements

We thank Dr. Mbaye Ndoye Sall (ISRA/CERAAS, Thies, Senegal) for kindly sharing the GBS data for the sorghum SSD population, Karine Labadie (CEA, Institut de Génomique, Genoscope, Evry, France) for sharing the WGS data for the rice RIL population, Christine Tranchant (IRD, Montpellier, France) for her help with retrieving the Rice_WGS data and François Sabot (IRD, Montpellier, France) for coordinating the IRIGIN project.

## Author’s contributions

ML developed the imputation algorithm and code for NOISYmputer, and wrote the draft manuscript. AG designed the bioinformatics pipeline to call SNPs for the Rice_WGS dataset. CF ran LB-Impute for the WGS_Rice dataset on the Yale cluster and edited the manuscript. JFR designed the pipelines for the Sorghum_GBS dataset and early tests of ABHgenotypeR.

## Notes

http://mapdisto.free.fr/Resources/files/NOISYmputer_Supplementary_Materials.zip

## References

Broman, K. W., Wu, H., Sen, Ś., & Churchill, G. A. (2003). R/qtl: QTL mapping in experimental crosses. Bioinformatics. https://doi.org/10.1093/bioinformatics/btg112

Davey, J. L., & Blaxter, M. W. (2010). RADseq: Next-generation population genetics. Briefings in Functional Genomics. https://doi.org/10.1093/bfgp/elq031

Elshire, R. J., Glaubitz, J. C., Sun, Q., Poland, J. A., Kawamoto, K., Buckler, E. S., & Mitchell, S. E. (2011). A robust, simple genotyping-by-sequencing (GBS) approach for high diversity species. PloS One. https://doi.org/10.1371/journal.pone.0019379

Fragoso, C. A., Heffelfinger, C., Zhao, H., & Dellaporta, S. L. (2016). Imputing genotypes in biallelic populations from low-coverage sequence data. Genetics. https://doi.org/10.1534/genetics.115.182071

Fragoso, C. A., Moreno, M., Wang, Z., Heffelfinger, C., Arbelaez, L. J., Aguirre, J. A., … Lorieux, M. (2017). Genetic Architecture of a Rice Nested Association Mapping Population. G3: Genes|Genomes|Genetics. https://doi.org/10.1534/g3.117.041608

Furuta, T., Ashikari, M., Jena, K. K., Doi, K., & Reuscher, S. (2017). Adapting Genotyping-by-Sequencing for Rice F2 Populations. G3: Genes|Genomes|Genetics. https://doi.org/10.1534/g3.116.038190

Gonen, S., Wimmer, V., Gaynor, R. C., Byrne, E., Gorjanc, G., & Hickey, J. M. (2018). A heuristic method for fast and accurate phasing and imputation of single-nucleotide polymorphism data in bi-parental plant populations. Theoretical and Applied Genetics. https://doi.org/10.1007/s00122-018-3156-9

Heffelfinger, C., Fragoso, C. A., & Lorieux, M. (2017). Constructing linkage maps in the genomics era with MapDisto 2.0. Bioinformatics. https://doi.org/10.1093/bioinformatics/btx177

Heffelfinger, C., Fragoso, C. A., Moreno, M. A., Overton, J. D., Mottinger, J. P., Zhao, H., … Dellaporta, S. L. (2014). Flexible and scalable genotyping-by-sequencing strategies for population studies. BMC Genomics. https://doi.org/10.1186/1471-2164-15-979

Hickey, J. M., Gorjanc, G., Varshney, R. K., & Nettelblad, C. (2015). Imputation of single nucleotide polymorphism genotypes in biparental, backcross, and topcross populations with a hidden markov model. Crop Science. https://doi.org/10.2135/cropsci2014.09.0648

Huang, X., Feng, Q., Qian, Q., Zhao, Q., Wang, L., Wang, A., … Han, B. (2009). High-throughput genotyping by whole-genome resequencing. Genome Research. https://doi.org/10.1101/gr.089516.108

Lander, E. S., Green, P., Abrahamson, J., Barlow, A., Daly, M. J., Lincoln, S. E., & Newburg, L. (1987). MAPMAKER: An interactive computer package for constructing primary genetic linkage maps of experimental and natural populations. Genomics. https://doi.org/10.1016/0888-7543(87)90010-3

Mace, E. S., Rami, J. F., Bouchet, S., Klein, P. E., Klein, R. R., Kilian, A., … Jordan, D. R. (2009). A consensus genetic map of sorghum that integrates multiple component maps and high-throughput Diversity Array Technology (DArT) markers. BMC Plant Biology. https://doi.org/10.1186/1471-2229-9-13

Miao, C., Fang, J., Li, D., Liang, P., Zhang, X., Yang, J., … Tang, H. (2018). Genotype-Corrector: Improved genotype calls for genetic mapping in F2 and RIL populations. Scientific Reports. https://doi.org/10.1038/s41598-018-28294-0

Swarts, K., Li, H., Romero Navarro, J. A., An, D., Romay, M. C., Hearne, S., … Bradbury, P. J. (2015). Novel Methods to Optimize Genotypic Imputation for Low-Coverage, Next-Generation Sequence Data in Crop Plants. The Plant Genome. https://doi.org/10.3835/plantgenome2014.05.0023

