## Supplementary material for "NOISYmputer: genotype imputation in bi-parental populations for noisy low-coverage next-generation sequencing data": NOISYmputer - Supplementary Data.docx

^*^Corresponding author

### GBS, WGS and bioinformatics pipelines

**Table S1.** Summary of methods used to produce real datasets

| ***Dataset*** | ***Cross*** | ***Pop. Type*** | ***Pop. size*** | ***Sequencing method*** | ***Sequencing technology*** | ***Sequencing depth*** | ***Data treatment pipeline*** | ***Pre-filtering*** |
| --- | --- | --- | --- | --- | --- | --- | --- | --- |
| Rice_GBS | [IR64 x Azucena] | SSD | 187 | GBS – Yale protocol [Heffelfinger et al 2014] – enzyme: RsaI | Illumina HiSeq 2500, paired-end 2 x 150 | ~0.67 X | Reads mapping: NovoAlign - SNP calling: GATK |  |
| Rice_WGS | [IR64 x Azucena] | SSD | 212 | WGS – Genoscope protocol [pers. comm] | Illumina HiSeq 4000 paired-end 2 x 150 | ~1.55 X | Reads mapping: BWA - SNP calling: GATK |  |
| Sorghum_GBS | [SSM1611 x SSM249] | SSD | 151 | GBS – Cornell [Elshire et al 2011] – enzyme: ApeKI | Illumina HiSeq 3000 paired-end 2 x 150 | ~0.77 X | Reads mapping: Bowtie2.3.2 - SNP calling: Tassel-GBS2 | MAF = 0.05 – Max missing data = 0.8 |

##### Rice_GBS

Details of GBS data preparation are in the Nested Association Mapping publication [Fragoso et al 2017]. Briefly, GBS paired-end reads of each SSD and parental line were aligned with Novoalign to the Nipponbare v7 reference genome. Variant calling was performed with GATK and a custom script described in the GBS publication [Heffelfinger et al 2014] was used for variant filtering. Each RIL population from the 2017 publication, including the IR64 x Azucena RIL advanced to F_10 or_ F_11_ via single seed descent (the dataset used in this current publication), was treated separately for variant calling and filtering.

##### Rice_WGS

Custom WGS libraries were prepared at Genoscope (Karine Labadie, pers. comm.). Paired-end reads were aligned with BWA to the Azucena reference genome (Rod Wing, pers. comm.). Variant calling was performed with GATK. Parental lines, which were sequenced at ~40 X, were treated separately from the SSD population, and the intersection of SNPs found between the parents and within the parents were retained. A custom Java program was then used for variant filtering.

##### Sorghum_GBS

GBS libraries were prepared using the original GBS protocol [Elshire et al 2011]. The Tassel GBS pipeline v. 2 was used using the following commands:

qsub -R yes -cwd -q normal.q -V -l mem_free=80G -b yes -N GBSSeqToTagDBPlugin "run_pipeline.pl -Xms10G -Xmx48G -fork1 -GBSSeqToTagDBPlugin -e ApeKI -i 01_RawSequence/ -db OutputTassel/GBSMb.db -k 50_KeyFiles/keyfile_tass.txt -mnQS 20 -kmerLength 64 -mxKmerNum 1000000000 -batchSize 653 -endPlugin -runfork1"

qsub -hold_jid GBSSeqToTagDBPlugin -R yes -N TagExportToFastqPlugin -cwd -q normal.q -V -l mem_free=80G -b yes "run_pipeline.pl -Xms10G -Xmx48G -fork1 -TagExportToFastqPlugin -db OutputTassel/GBSMb.db -o 02_Tag_fastq/GBSMb.fastq -c 5 -endPlugin -runfork1"

qsub -hold_jid TagExportToFastqPlugin -V -q normal.q -b y -N Aln "bowtie2 -p 1 --very-sensitive-local -q 02_Tag_fastq/GBSMb.fastq -x /bank/sorghum_bicolor/v3.1/Sbicolor_313_v3.0.fa -S 02_Tag_fastq/GBSMb.sam"

qsub -hold_jid Aln -N bowtie2_clean -cwd -q normal.q -V -l mem_free=80G -b yes "perl utils/filter_sam_file_GBS.pl 02_Tag_fastq/GBSMb.sam"

qsub -hold_jid bowtie2_clean -N SAMToGBSdbPlugin -cwd -q normal.q -V -l mem_free=80G -b yes "run_pipeline.pl -Xms10G -Xmx48G -fork1 -SAMToGBSdbPlugin -i 02_Tag_fastq/GBSMb.sam.clean -db OutputTassel/GBSMb.db -endPlugin -runfork1"

qsub -hold_jid SAMToGBSdbPlugin -N DiscoverySNPCallerPluginV2 -cwd -q normal.q -V -l mem_free=80G -b yes "run_pipeline.pl -Xms10G -Xmx48G -fork1 -DiscoverySNPCallerPluginV2 -db OutputTassel/GBSMb.db -mnLCov 0.1 -endPlugin -runfork1"

qsub -hold_jid DiscoverySNPCallerPluginV2 -N SNPQualityProfilerPlugin -cwd -q normal.q -V -l mem_free=80G -b yes "run_pipeline.pl -Xms10G -Xmx48G -fork1 -SNPQualityProfilerPlugin -db OutputTassel/GBSMb.db -tname GBSMb -statFile outputStats-GBSMb -endPlugin -runfork1"

qsub -hold_jid SNPQualityProfilerPlugin -N ProductionSNPCallerPluginV2 -cwd -q normal.q -V -l mem_free=80G -b yes "run_pipeline.pl -Xms10G -Xmx48G -fork1 -ProductionSNPCallerPluginV2 -db OutputTassel/GBSMb.db -e ApeKI -eR 0.01 -i 01_RawSequence/ -k 50_KeyFiles/keyfile_tass.txt -o 03_Production_SNP_Caller/GBSMb.h5 -mnQS 20 -kmerLength 64 -endPlugin -runfork1"

qsub -hold_jid ProductionSNPCallerPluginV2 -N exportVCF -cwd -q normal.q -V -l mem_free=80G -b yes "run_pipeline.pl -Xms10G -Xmx48G -h5 03_Production_SNP_Caller/GBSMb.h5 -export 03_Production_SNP_Caller/GBSMb.vcf -exportType VCF"

### Imputation commands and parameters

#### NOISYmputer

##### Rice_GBS datasets

**Pre-filtering**

max missing data = 0.8, max heterozygosity = 0.025, min f(A) = 0.05, min f(B) = 0.05

**Filtering incoherent SNPs**

½ window size = 30, shift = 1, Chi^2^ threshold = 3.84

**Imputation step 1**

½ window size = 30, min f(A) in A = 0.9, min f(B) in B = 0.95, Htz transition rate = 0.2, min f(A) in H = 0.1, min f(B) in H = 0.05

**Imputation step 2**

½ window size = 25, min f(A) = 0.6, min f(B) = 0.7, min f(H) = 0.15, min f(A) in H = 0.33, min f(B) in H = 0.55

**Imputation step 3**

½ window size = 15

**Improbable chunk detection**

Max chunk size = 35, Chunk environment = 0.5, method = 2, min local recomb rate = 15, max local recomb rate = 50

##### Sorghum_GBS datasets

**Pre-filtering**

max missing data = 0.8, max heterozygosity = 0.025, min f(A) = 0.05, min f(B) = 0.05

**Filtering incoherent SNPs**

½ window size = 30, shift = 1, Chi^2^ threshold = 3.84

**Imputation step 1**

½ window size = 30, min f(A) in A = 0.9, min f(B) in B = 0.9, Htz transition rate = 0.2, min f(A) in H = 0.1, min f(B) in H = 0.05

**Imputation step 2**

½ window size = 25, min f(A) = 0.6, min f(B) = 0.6, min f(H) = 0.15, min f(A) in H = 0.55, min f(B) in H = 0.55

**Imputation step 3**

½ window size = 15

**Improbable chunk detection**

Max chunk size = 35, Chunk environment = 0.5, method = 2, min local recomb rate = 15, max local recomb rate = 50

##### Rice_WGS dataset

**Pre-filtering**

max missing data = 0.6, max heterozygosity = 0.025, min f(A) = 0.01, min f(B) = 0.01

**Filtering incoherent SNPs**

½ window size = 30, shift = 1, Chi^2^ threshold = 3.84

**Imputation step 1**

½ window size = 30, min f(A) in A = 0.9, min f(B) in B = 0.95, Htz transition rate = 0.2, min f(A) in H = 0.1, min f(B) in H = 0.05

**Imputation step 2**

½ window size = 25, min f(A) = 0.6, min f(B) = 0.75, min f(H) = 0.075, min f(A) in H = 0.33, min f(B) in H = 0.55

**Imputation step 3**

½ window size = 15

**Improbable chunk detection**

Max chunk size = 500, Chunk environment = 0.5, method = 2, min local recomb rate = 15, max local recomb rate = 50

#### LB-Impute

##### Rice_GBS and Sorghum_GBS datasets

**Pre-filtering with Tassel**

Site Min Count = 50, Site Min Allele Freq = 0.2, Site Max Allele Freq = 0.8, Min Heterozygous Proportion = 0, Max Heterozygous Proportion = 0.05, all other parameters left as default value

**Parental imputation**

java -jar LB-Impute_v1.jar -method impute -f VCFname.vcf -parents Parent1,Parent2 -o VCFname_imputed_Parents.vcf -window 7 -minsamples 5 -minfraction 0.5 -recombdist 10000000 -genotypeerr 0.05 -readerr 0.05 -dr -parentimpute

**Offspring imputation**

java -jar LB-Impute_v1.jar -method impute -f VCFname_imputed_Parents.vcf -parents Parent1,Parent2 -o VCFname_imputed_Offspring.vcf -window 7 -minsamples 5 -minfraction 0.5 -recombdist 10000000 -genotypeerr 0.05 -readerr 0.05 -dr -offspringimpute

##### Rice_WGS dataset

**VCF formatting**

The following Linux sed commands were used to convert vcf 4.2 format into a format readable by LB-Impute. These commands remove the allele balance “AB” format field from the vcf file.

sed -i 's/GT:AB:/GT:/g' VCFname.vcf;

sed -i 's/0\/0:\.:/0\/0:/g' VCFname.vcf;

sed -i 's/1\/1:\.:/1\/1:/g' VCFname.vcf ;

sed -i 's/0\/1:[0-9]\.[0-9][0-9][0-9]:/0\/1:/g' VCFname.vcf;

sed -i 's/0\/1:0\.00:/0\/1:/g' VCFname.vcf;

sed -i 's/0\/1:1\.00:/0\/1:/g' VCFname.vcf;

**Parental imputation**

java -jar LB-Impute_v1.jar -method impute -f VCFname.vcf -parents Parent1,Parent2 -o VCFname_imputed_Parents.vcf -window 7 -minsamples 5 -minfraction 0.5 -recombdist 10000000 -genotypeerr 0.05 -readerr 0.05 -dr -parentimpute

**Offspring imputation**

java -jar LB-Impute_v1.jar -method impute -f VCFname_imputed_Parents.vcf -parents Parent1,Parent2 -o VCFname_imputed_Offspring.vcf -window 7 -minsamples 5 -minfraction 0.5 -recombdist 10000000 -genotypeerr 0.05 -readerr 0.05 -dr -offspringimpute

#### FSFHap (Tassel 5, GUI, macOS)

##### Rice_GBS dataset

**Pre-filtering**

Site Min Count = 50, Site Min Allele Freq = 0.1, Site Max Allele Freq = 0.9, Min Heterozygous Proportion = 0, Max Heterozygous Proportion = 0.05, all other parameters left as default value

**Imputation**

Use Cluster Algorithm = true, Min Minor Allele Frequency = 0.1, Window Size = 50, Min R = 0.2, Max Missing = 0.8, Don’t Use Heterozygous Calls = false, Max Diff = 0, Min Hap = 2, Window Overlap = 25, Fill Gaps = false, Proportion Heterozygous = 0.02, Merge = false, Out Parents = true, Out Nucleotides = true, Output IUPAC Codes = true

##### Sorghum_GBS dataset

**Pre-filtering**

Site Min Count = 50, Site Min Allele Freq = 0.2, Site Max Allele Freq = 0.8, Min Heterozygous Proportion = 0, Max Heterozygous Proportion = 0.05, all other parameters left as default value

**Imputation**

Use Cluster Algorithm = true, Min Minor Allele Frequency = 0.1, Window Size = 50, Min R = 0.2, Max Missing = 0.8, Don’t Use Heterozygous Calls = false, Max Diff = 0, Min Hap = 2, Window Overlap = 25, Fill Gaps = false, Proportion Heterozygous = 0.02, Merge = false, Out Parents = true, Out Nucleotides = true, Output IUPAC Codes = true

##### Rice_WGS dataset

**Pre-filtering**

Site Min Count = 50, Site Min Allele Freq = 0.1, Site Max Allele Freq = 0.9, Min Heterozygous Proportion = 0, Max Heterozygous Proportion = 0.05, all other parameters left as default value

**Imputation**

Use Single Backcross Algorithm = true^*^, Min Minor Allele Frequency = 0.1, Window Size = 50, Min R = 0.2, Max Missing = 0.8, Don’t Use Heterozygous Calls = false, Max Diff = 0, Min Hap = 2, Window Overlap = 25, Fill Gaps = false, Proportion Heterozygous = 0.02, Merge = false, Out Parents = true, Out Nucleotides = true, Output IUPAC Codes = true

^*^FSFHap generated an error with the Cluster Algorithm, so we used the Single Backcross algorithm instead

#### Genotype-Corrector

**VCF to .map conversion**

We used the NOISYmputer ‘Convert VCF to ABH genotypes’ function, as the vcf2map command in Genotype-Corrector cannot use parental genotypes to phase the data

**Filter out missing data**

python3 -m schnablelab.imputation.GC qc_missing NGS_Dataset_filtered.map NGS_Dataset _QCmissing.map

**Filter out segregation distortion**

python3 -m schnablelab.imputation.GC qc_sd NGS_Dataset_QCmissing.map NGS_Dataset_QCsd.map --population RIL

**Merge the consecutive same homozygous markers within a short genomic interval in the heterozygous region**

python3 -m schnablelab.imputation.GC qc_hetero NGS_Dataset_QCsd.map NGS_Dataset_QChetero.map

**Run correction**

python3 -m schnablelab.imputation.GC correct GCconf.txt NGS_Dataset_QChetero.map --debug

Configuration file:

[Section1]

Population_type: RIL

[Section2]

Letter_for_homo1: A

Letter_for_hete: X

Letter_for_homo2: B

Letter_for_missing_data: -

[Section3]

error_rate_for_homo1: 0.03

error_rate_for_homo2: 0.01

[Section4]

Sliding_window_size: 15 (1135 for the Rice_WGS dataset)

**Bin consecutive markers with same genotypes**

python3 -m schnablelab.imputation.GC bin NGS_Dataset_QChetero.corrected.map NGS_Dataset_QChetero.final.map

### Imputation results (graphical genotypes)

#### Rice_GBS dataset

**Figure S1.** Partial graphical genotypes plot of Rice_GBS high-quality data (chromosome 1), imputed with **NOISYmputer**. SSD lines go in columns and SNPs go in rows. Blue: Azucena allele; red: IR64 allele; yellow: heterozygote; white: unimputed data

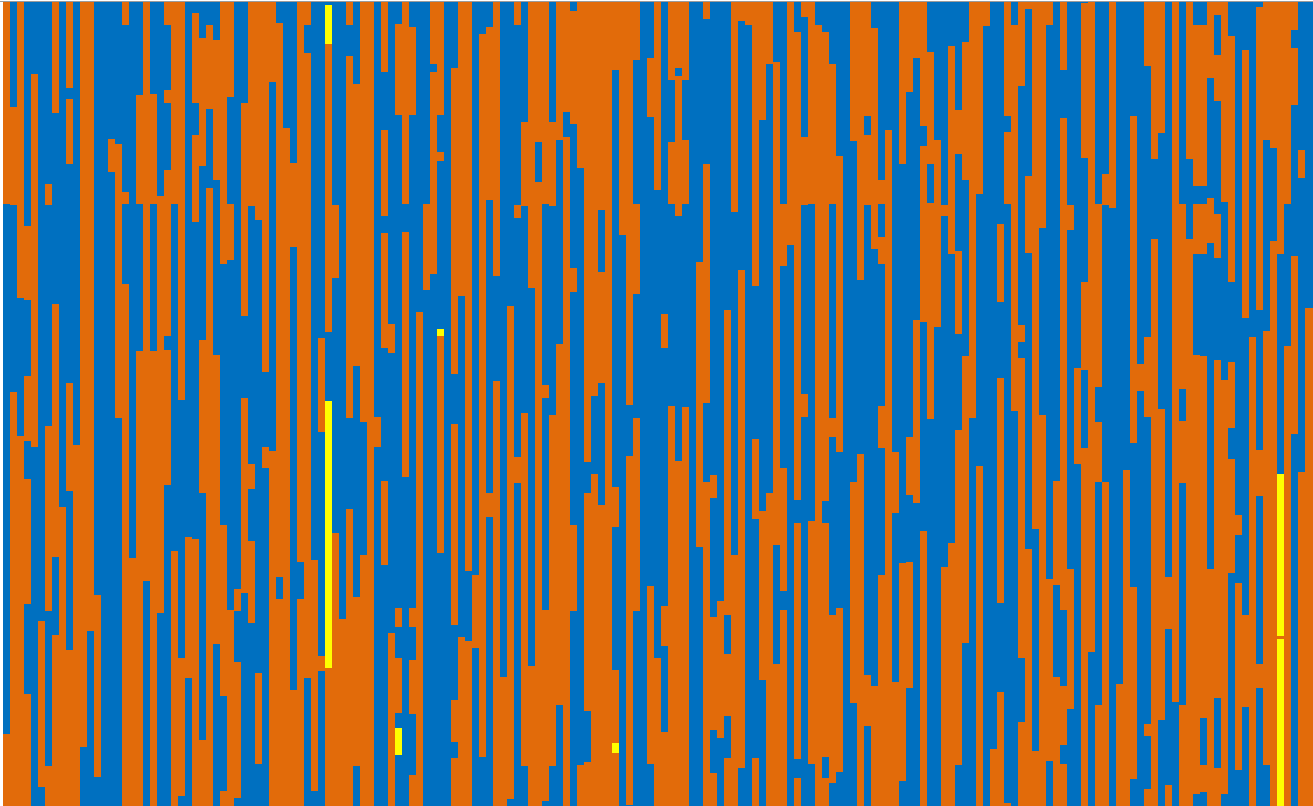

**Figure S2.** Partial graphical genotypes plot of Rice_GBS high-quality data (chromosome 1), imputed with **LB-Impute**. SSD lines go in columns and SNPs go in rows. Blue: Azucena allele; red: IR64 allele; yellow: heterozygote; white: unimputed data

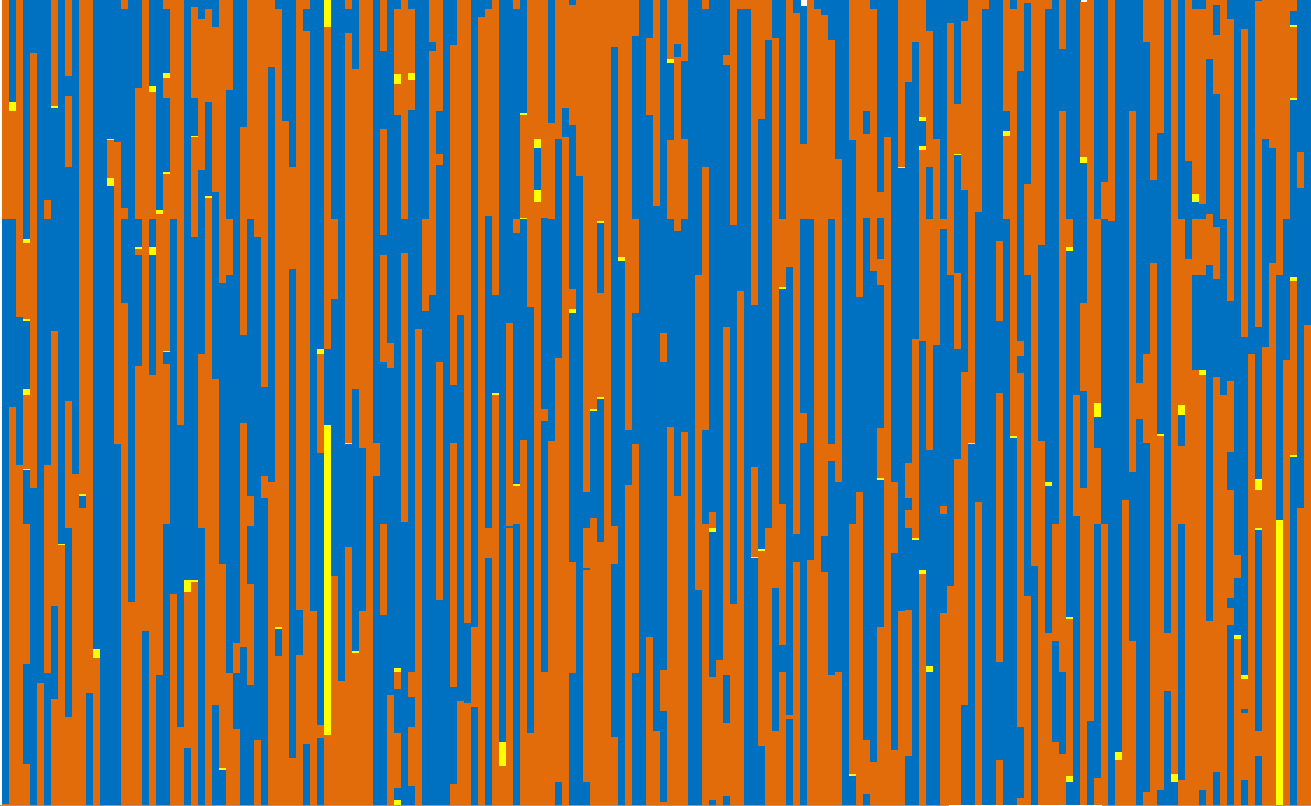

**Figure S3.** Partial graphical genotypes plot of Rice_GBS high-quality data (chromosome 1), imputed with **FSFHap**. SSD lines go in columns and SNPs go in rows. Blue: Azucena allele; red: IR64 allele; yellow: heterozygote; white: unimputed data

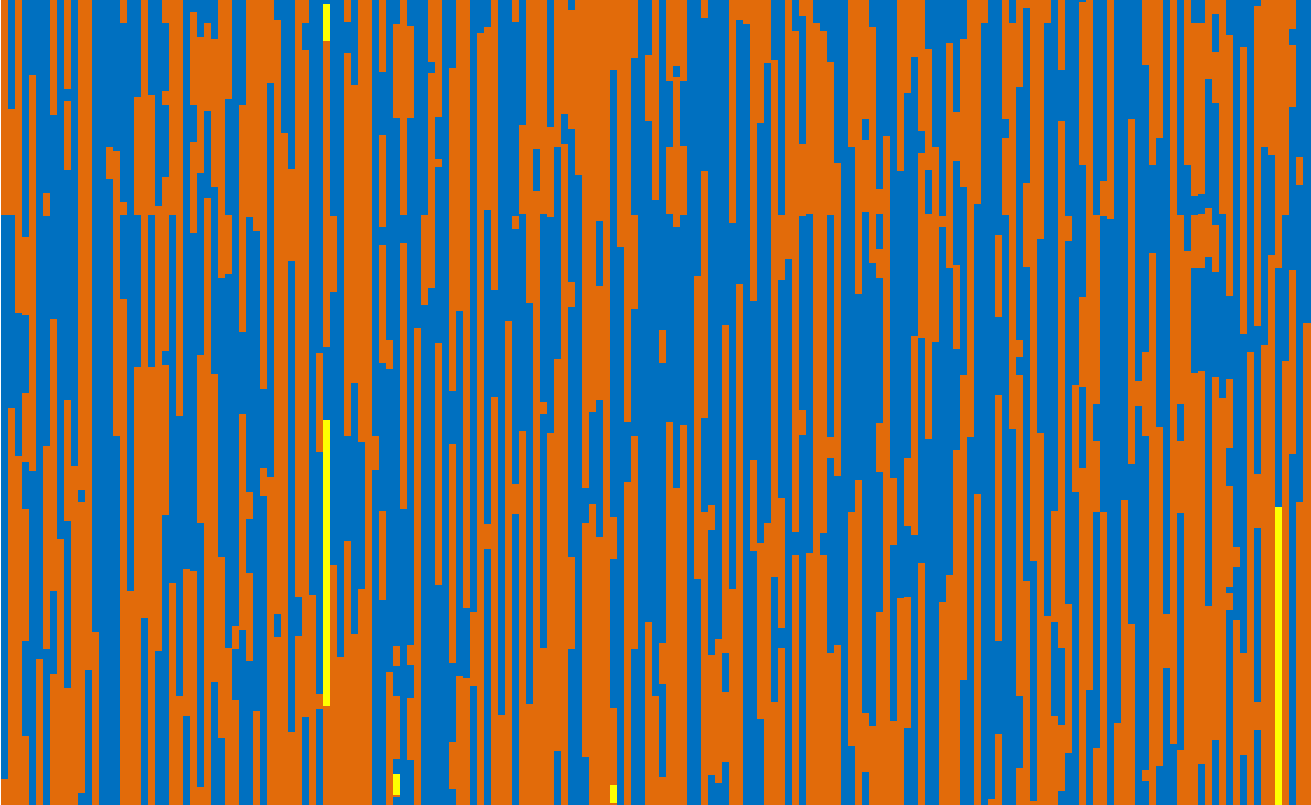

**Figure S4.** Partial graphical genotypes plot of Rice_GBS high-quality data (chromosome 1), imputed with **Genotype-Corrector**. SSD lines go in columns and SNPs go in rows. Blue: Azucena allele; red: IR64 allele; yellow: heterozygote; white: unimputed data

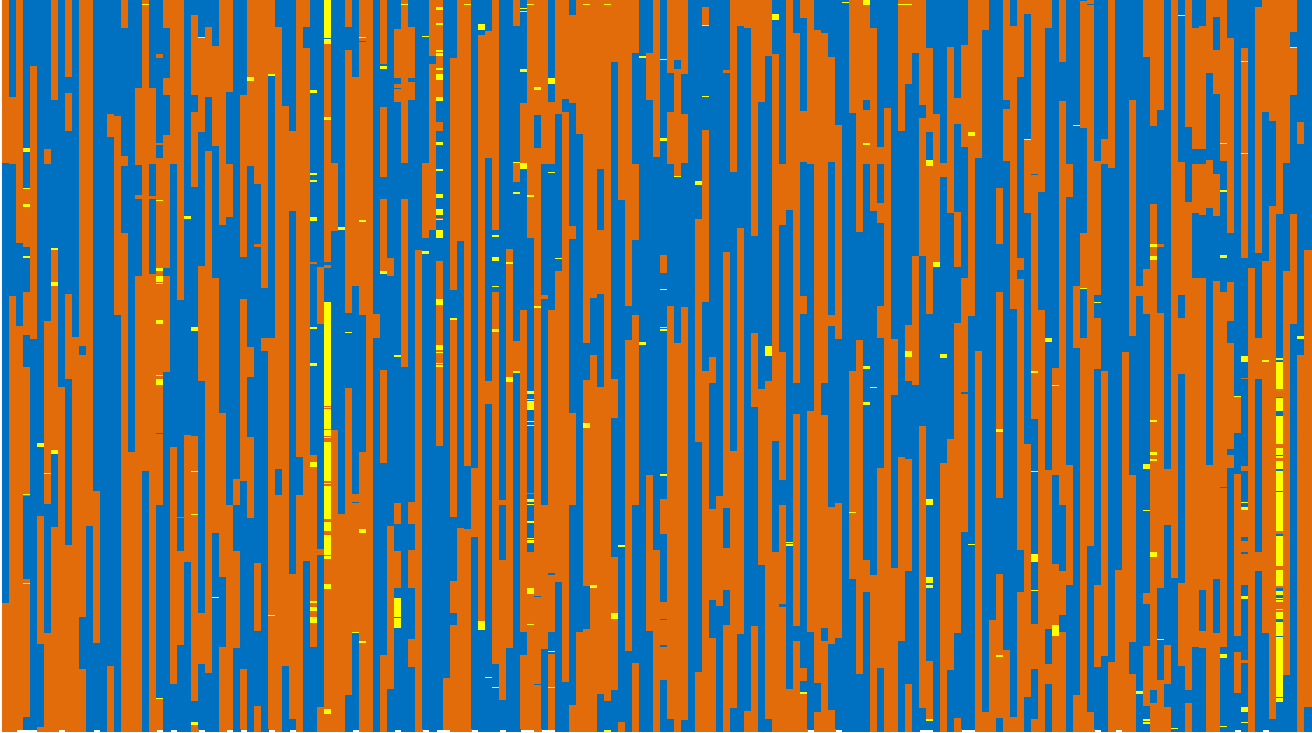

#### Sorghum_GBS dataset

**Figure S5.** Partial graphical genotypes plot of Sorghum_GBS noisy data (chromosome 1), imputed with **NOISYmputer**. SSD lines go in columns and SNPs go in rows. Blue: Azucena allele; red: IR64 allele; yellow: heterozygote; white: unimputed data. NOISYmputer filtered out all noisy data, resulting in correct final map size

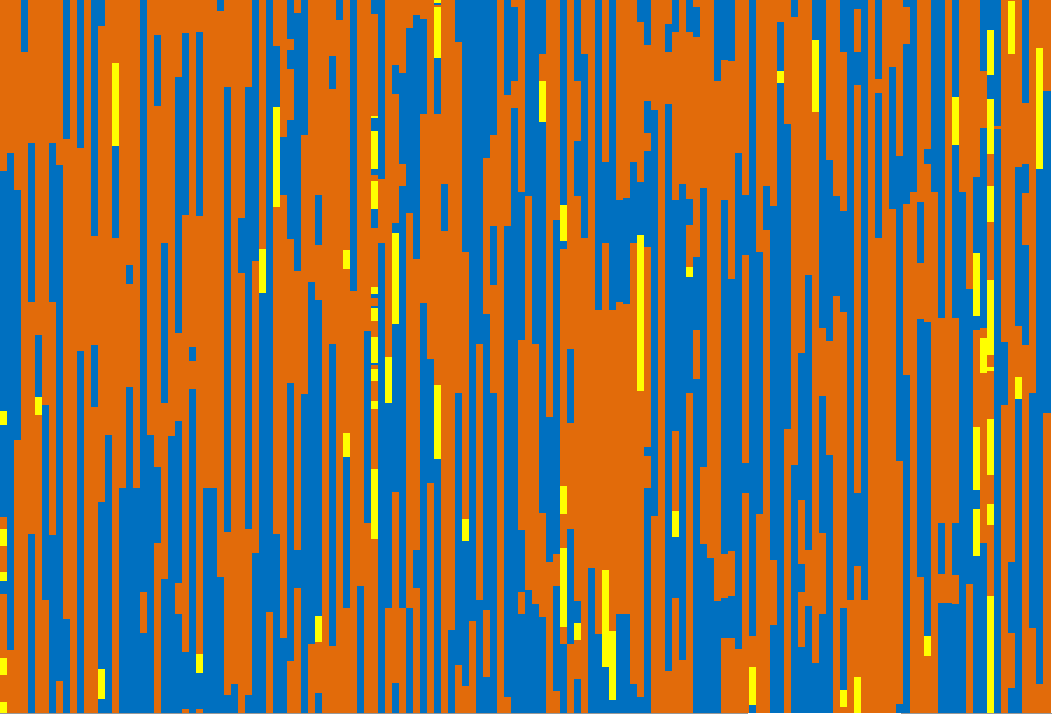

**Figure S6.** Partial graphical genotypes plot of Sorghum_GBS noisy data (chromosome 1), imputed with **LB-Impute**. SSD lines go in columns and SNPs go in rows. Blue: SSM1611 allele; red: SSM249 allele; yellow: heterozygote; white: unimputed data. LB-Impute inaccurately imputed many singletons, resulting in very severe map expansion (~53.4 X)

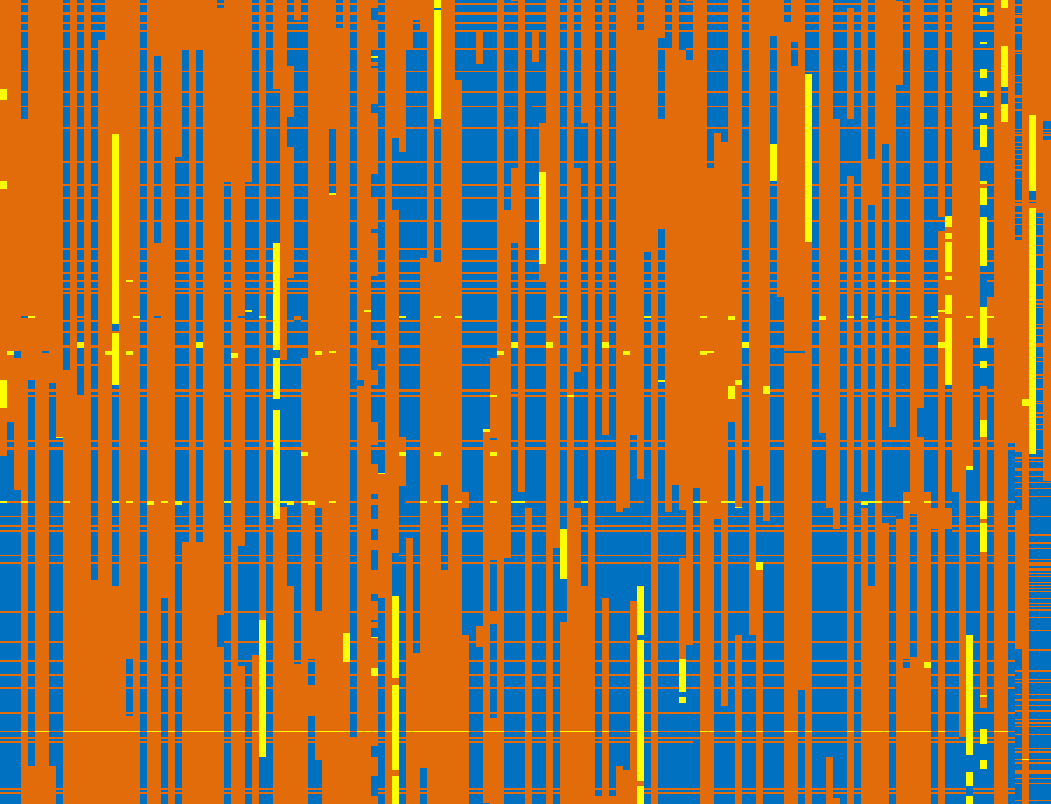

**Figure S7.** Partial graphical genotypes plot of Sorghum_GBS noisy data (chromosome 1), imputed with **FSFHap**. SSD lines go in columns and SNPs go in rows. Blue: SSM1611 allele; red: SSM249 allele; yellow: heterozygote; white: unimputed data. FSFHap inaccurately imputed many singletons, resulting in severe map expansion (~5.8 X)

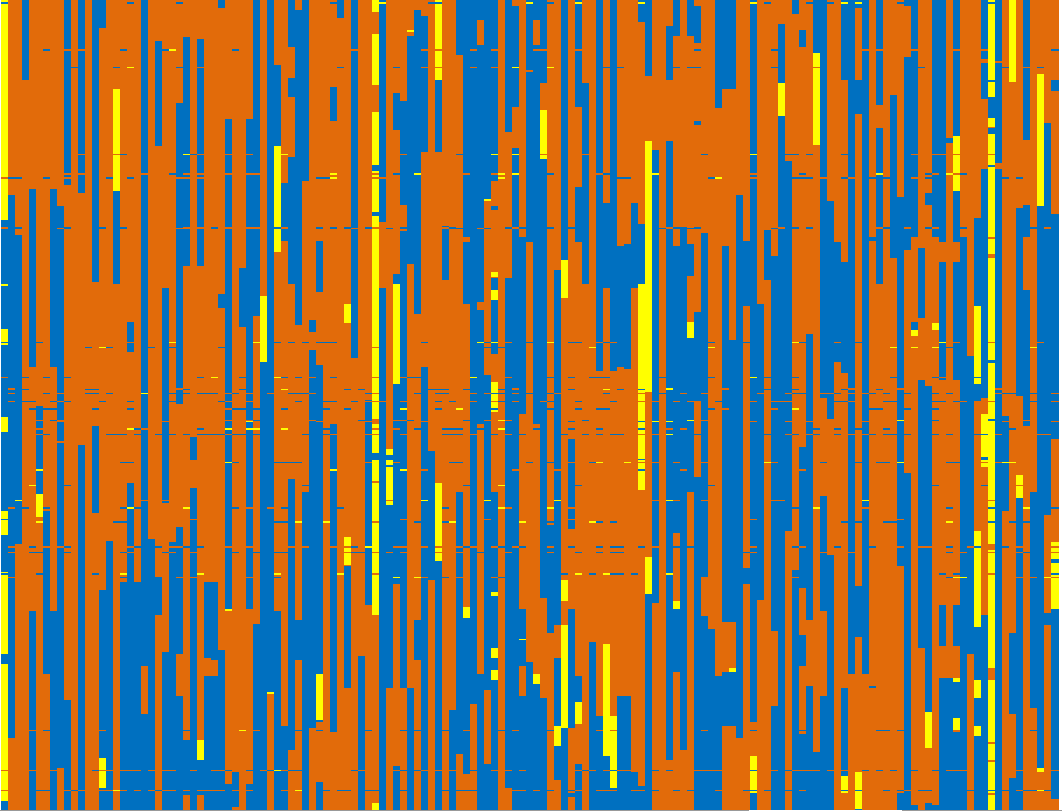

**Figure S8.** Partial graphical genotypes plot of Sorghum_GBS noisy data (chromosome 1), imputed with **Genotype-Corrector**. SSD lines go in columns and SNPs go in rows. Blue: SSM1611 allele; red: SSM249 allele; yellow: heterozygote; white: unimputed data. Genotype-Corrector was unable to eliminate some noisy data and converted them to heterozygous calls (particularly obvious for the range of SNPs approximately in the middle of the graph), resulting in severe map expansion (~3.7 X)

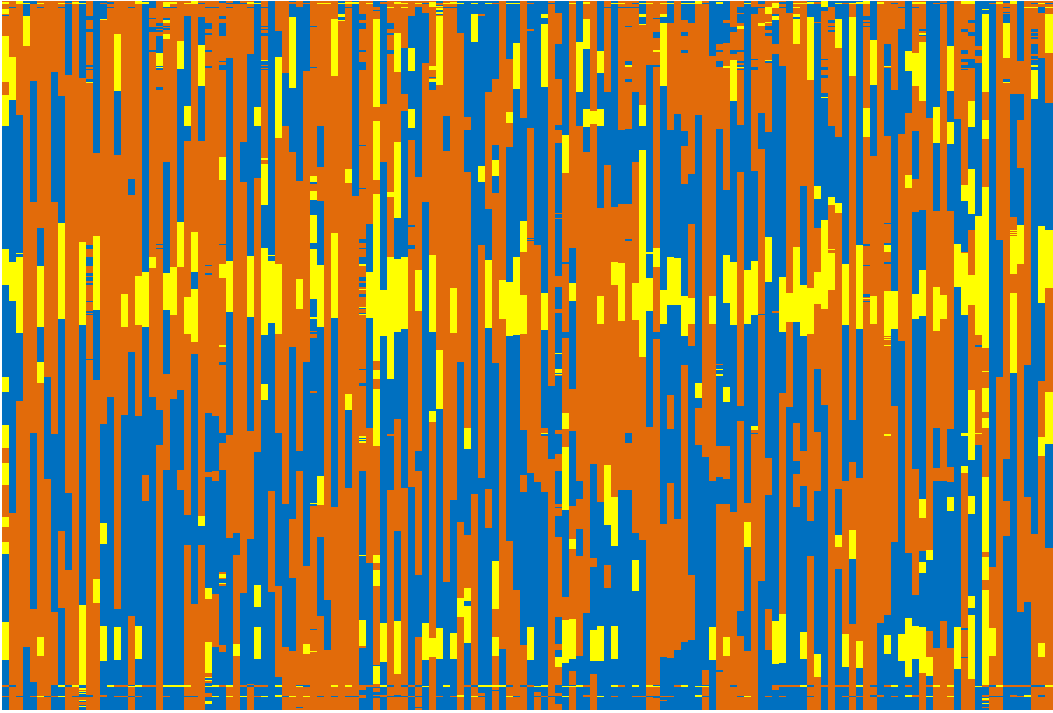

#### Rice_WGS dataset

**Figure S9.** Partial graphical genotypes plot of Rice_WGS noisy data (chromosome 1), imputed with **NOISYmputer**. SSD lines go in columns and SNPs go in rows. Blue: Azucena allele; red: IR64 allele; yellow: heterozygote; white: unimputed data

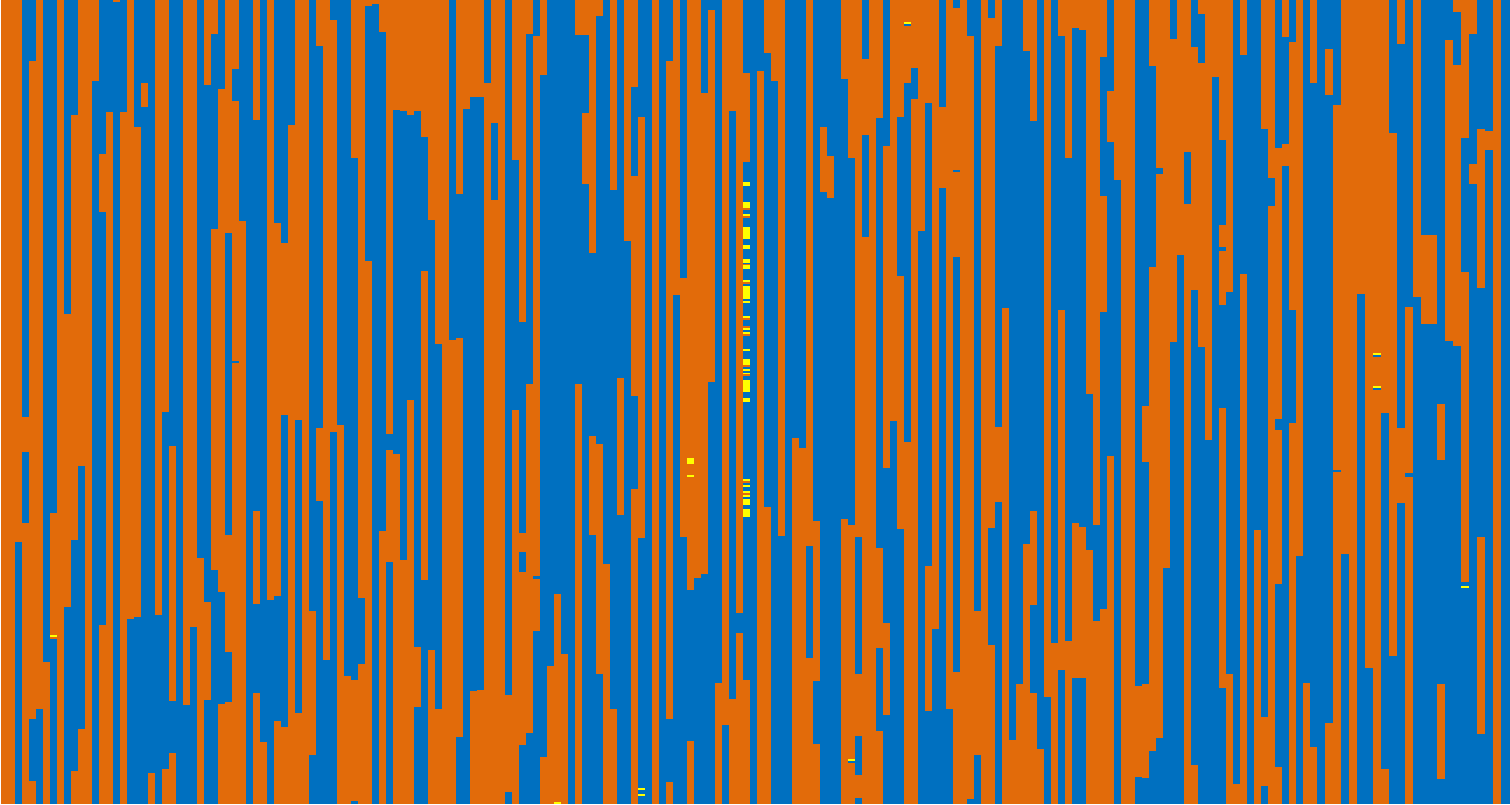

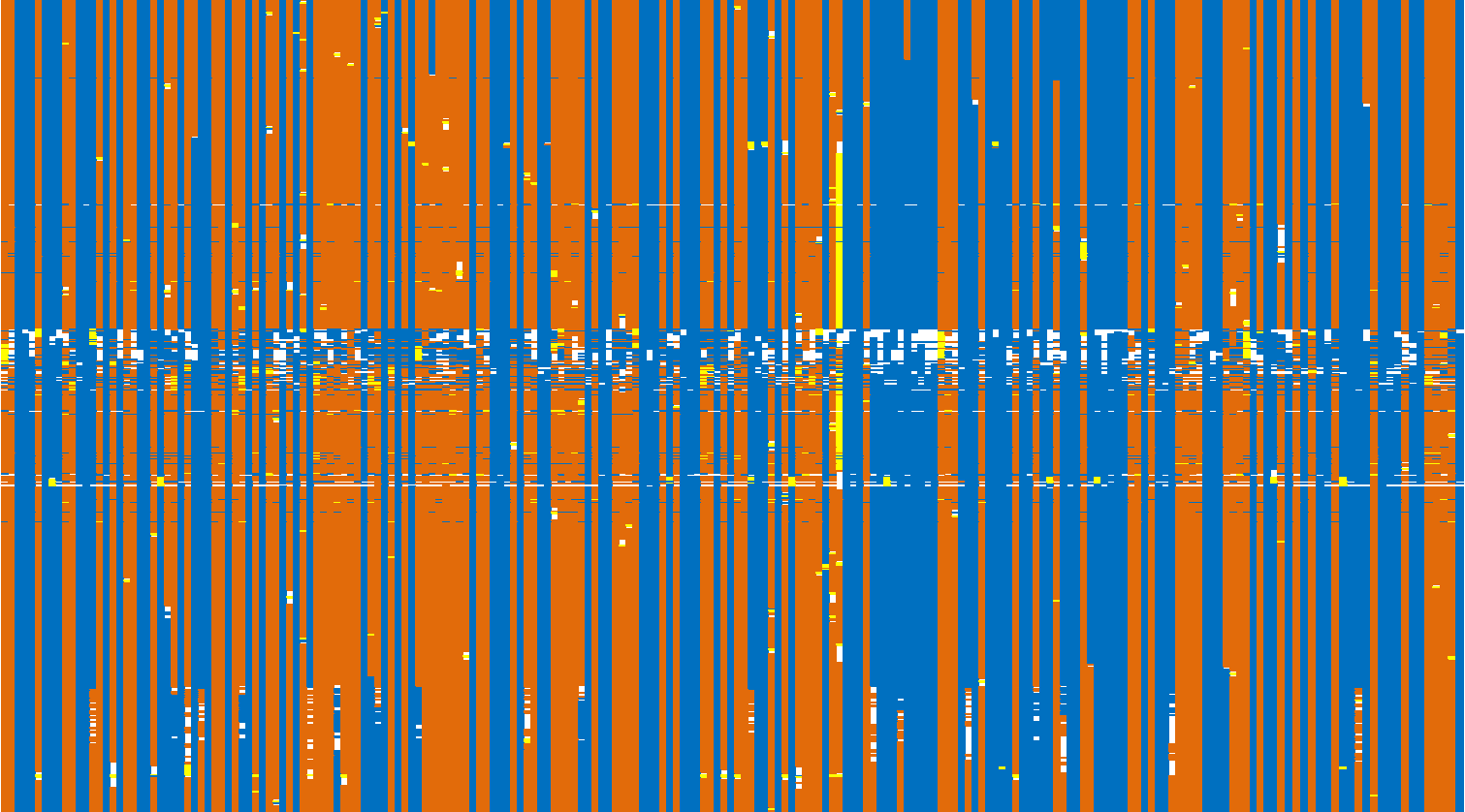
**Figure S10.** Partial graphical genotypes plot of Rice_WGS noisy data (chromosome 1), imputed with **FSFHap**. SSD lines go in columns and SNPs go in rows. Blue: Azucena allele; red: IR64 allele; yellow: heterozygote; white: unimputed data. The inability of FSFHap to eliminate noisy data (particularly obvious for the range of SNPs approximately in the middle of the graph), and the inaccurate imputation of many small heterozygous chromosome chunks conducted to severe map expansion

**Figure S11.** Partial graphical genotypes plot of Rice_WGS noisy data (chromosome 1), imputed with **Genotype-Corrector**. SSD lines go in columns and SNPs go in rows. Blue: Azucena allele; red: IR64 allele; yellow: heterozygote; white: unimputed data

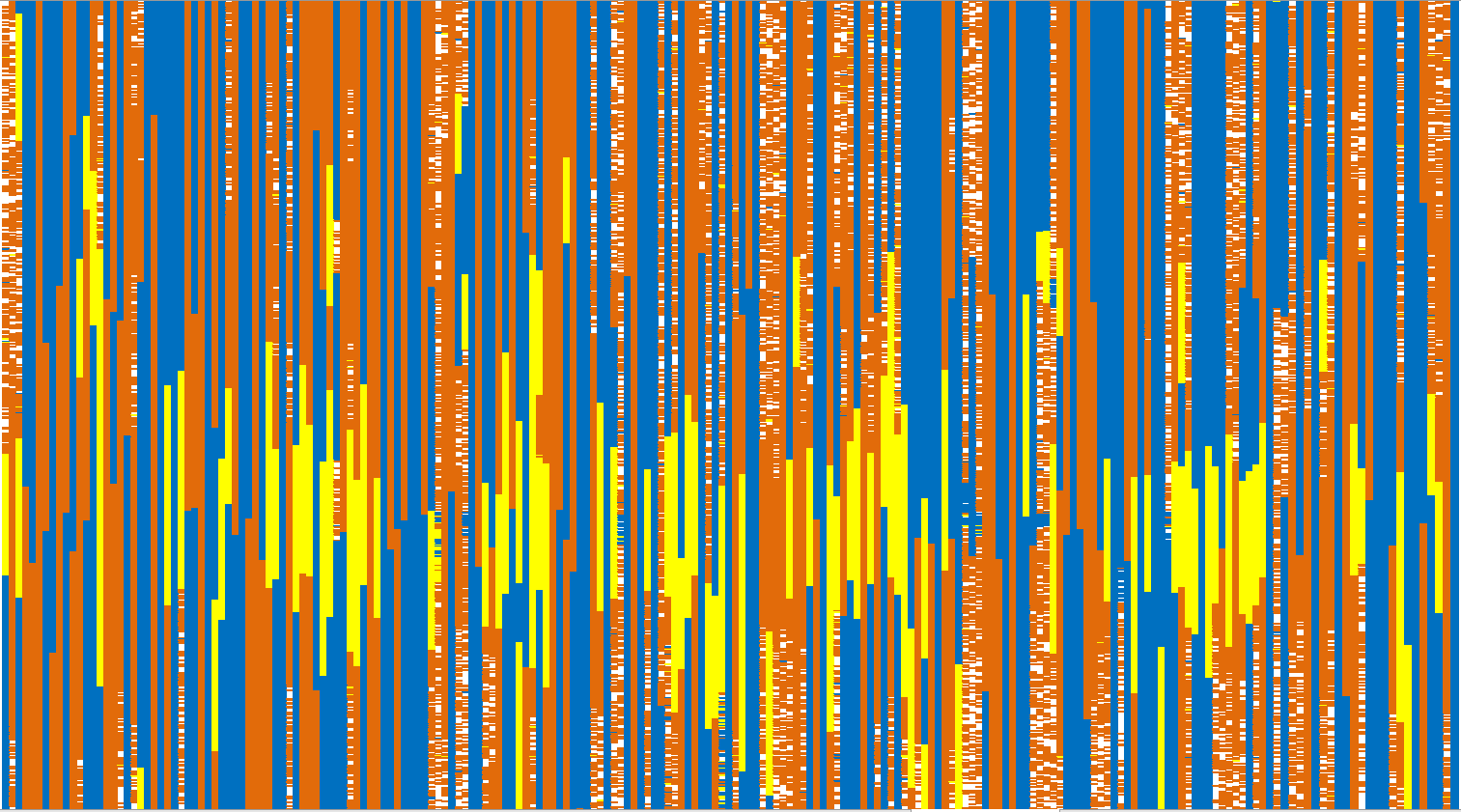

### References

Elshire, R. J., Glaubitz, J. C., Sun, Q., Poland, J. A., Kawamoto, K., Buckler, E. S., & Mitchell, S. E. (2011). A Robust, Simple Genotyping-by-Sequencing (GBS) Approach for High Diversity Species. *PLoS ONE*, *6*(5), e19379. http://doi.org/10.1371/journal.pone.0019379.g006

Fragoso, C. A., Moreno, M., Wang, Z., Heffelfinger, C., Arbelaez, L. J., Aguirre, J. A., et al. (2017). Genetic Architecture of a Rice Nested Association Mapping Population. *G3: Genes, Genomes, Genetics*, g3.117.041608–15. <http://doi.org/10.1534/g3.117.041608>

Heffelfinger, C., Fragoso, C. A., Moreno, M. A., Overton, J. D., Mottinger, J. P., Zhao, H., et al. (2014). Flexible and scalable genotyping-by-sequencing strategies for population studies. *BMC Genomics*, *15*(1), 979. <http://doi.org/10.1186/1471-2164-15-979>
